# GBactPro: cross-species bacterial promoter prediction using element-resolved promoter definitions

**DOI:** 10.64898/2026.10.18.759981

**Authors:** Hsiang-Yu Bruce Han, Yu-Chien Vicky Kao, Syue-Ting Antony Kuo, Hsin-Hung David Chou, Jia-Ming Chang

**Affiliations:** Department of Life Science, National Taiwan University, Taipei 106319, Taiwan; Department of Computer Science, National Chengchi University, Taipei 116011, Taiwan; Genome and Systems Biology Degree Program, Academia Sinica and National Taiwan University, Taipei 106319, Taiwan; Institute of Neuroscience, National Chengchi University, Taipei 116011, Taiwan

## Abstract

Accurate bacterial promoter prediction supports genome annotation, but many models use fixed-length promoter windows anchored to transcription start sites (TSSs) rather than promoter-element boundaries. We developed the <u>G</u>eneral <u>Bact</u>erial <u>Pro</u>moter Predictor (GBactPro), a convolutional neural network–bidirectional long short-term memory (CNN–BiLSTM) framework using promoter sequences resolved by the Promoter Architecture Scanner. Using *Escherichia coli*-trained models evaluated across 43 bacterial species, we compared eight input designs and identified the contiguous −35/spacer/−10 core as optimal (mean AUROC = 0.9859). Species-specific models showed broad transfer in a 43 × 43 evaluation. Joint training on all 43 species achieved a mean AUROC of 0.9943 across their held-out test sets and outperformed four established predictors under matched-length benchmarks. In genome-wide *E. coli* scanning, GBactPro achieved AUROC and AUPRC values of 0.979 and 0.160 under stringent full-containment matching (0-bp tolerance), increasing to 0.997 and 0.323 under a 100-bp tolerance. False-positive predictions overlapped H-NS binding regions more frequently than true positives (24.4% versus 18.8%), and predictions also overlapped promoters derepressed in an H-NS-knockout dataset. Prediction precision varied across four TSS classes. These findings demonstrate that element-resolved promoter definitions improve cross-species prediction and support genome-wide identification of candidate bacterial promoters.

## Introduction

Identifying bacterial promoters is essential for understanding transcriptional regulation and annotating transcriptional units and operons (1,2). Experimental approaches, including differential RNA sequencing (dRNA-seq), Cappable-seq, and SMRT-Cappable-seq, have enabled genome-wide mapping of transcription start sites (TSSs), providing extensive datasets for promoter characterization (3–5). However, experimentally mapped TSSs depend on the conditions examined and remain incomplete or unavailable for many bacterial species. Computational promoter prediction therefore provides a complementary approach for genome annotation and the identification of candidate regulatory regions (6,7). Importantly, promoter-like DNA sequences do not necessarily exhibit transcriptional activity in vivo. Recent experiments involving the deletion of nine abundant nucleoid-associated proteins (NAPs) in *Escherichia coli* revealed widespread transcriptional dysregulation and a more homogeneous distribution of transcriptional output across the genome (8).

Cross-species promoter prediction is particularly valuable for bacteria with limited transcriptomic data. The core architecture of bacterial σ70 promoters, comprising the −35 and −10 elements and their intervening spacer, is broadly conserved across diverse species, although the sequences and composition of these elements vary among bacterial lineages (9). These shared architectural features provide a biological basis for cross-species prediction. Nevertheless, models trained on promoter data from one species may exhibit reduced performance when applied to other species, highlighting the challenge of learning transferable promoter features (6,10–12).

An important consideration is how promoter sequences are defined for model training. Many supervised bacterial promoter predictors use fixed-length input windows positioned relative to annotated TSSs rather than constructing training examples from experimentally resolved promoter-element boundaries (11–16). Although computationally convenient, these windows may include sequence beyond the core promoter, including downstream transcribed regions in some designs, and may not consistently align the elements involved in promoter recognition. We therefore hypothesized that defining training sequences according to promoter architecture could improve cross-species prediction by emphasizing conserved core promoter features.

The Promoter Architecture Scanner (PAS), developed by Kuo et al. (9), provides a basis for investigating this hypothesis. By integrating high-throughput experimental data with biophysical modeling, PAS characterized promoter architecture upstream of experimentally mapped TSSs across 49 bacterial species, resolving the −35 and −10 elements and their intervening spacer together with other promoter-associated elements. These annotations enable promoter training sequences to be defined by their constituent elements rather than by predetermined window boundaries. Whereas PAS characterizes promoter architecture using known TSS positions, these annotations provide the foundation for training a predictor that identifies candidate promoter regions without requiring TSS annotations during inference.

Previous computational approaches have explored diverse sequence features and model architectures. Early methods employed neural networks (17), variable-window Z-curve representations (18), and conventional feature-based classifiers (13,19,20), whereas bTSSfinder incorporated promoter-element information for predicting different sigma-factor classes (21). Subsequent developments introduced convolutional neural networks (22), random-forest-based prediction in Promotech (11), ensemble learning in iPro-WAEL (14), Siamese neural networks in TIMER (23), and multilayer perceptron-based prediction in Prompt (15). More recently, pretrained language models have been applied to promoter prediction, including BERT-Promoter (24), ProkBERT (25), iProL (16), msBERT-Promoter (26), and DNABERT-based iPro-MP (12). PromoterAtlas further extended this direction by pretraining a transformer model on approximately 9 million regulatory sequences from 3,371 gammaproteobacterial species and using a segmentation-based framework to annotate promoter elements associated with five sigma-factor classes (27). Despite these advances in sequence representation and promoter-element annotation, the influence of experimentally resolved promoter-element boundaries on supervised training and cross-species prediction remains comparatively less explored.

Here, we developed the General Bacterial Promoter Predictor (GBactPro), a convolutional neural network– bidirectional long short-term memory (CNN–BiLSTM) framework, to investigate how promoter-sequence definition affects cross-species prediction (Fig. 1). We systematically compared eight input-sequence designs using models trained on *E. coli* and evaluated across 43 bacterial species, identifying the contiguous −35/spacer/−10 core as the optimal input. Using this input definition, we assessed cross-species transfer through a 43 × 43 evaluation of species-specific models, examined the benefits of joint multi-species training, and benchmarked GBactPro against four established predictors under matched-length settings. Finally, we applied GBactPro to genome-wide promoter-site prediction in *E. coli* and investigated its predictions in relation to H-NS binding and four TSS classes.

**Figure 1.**
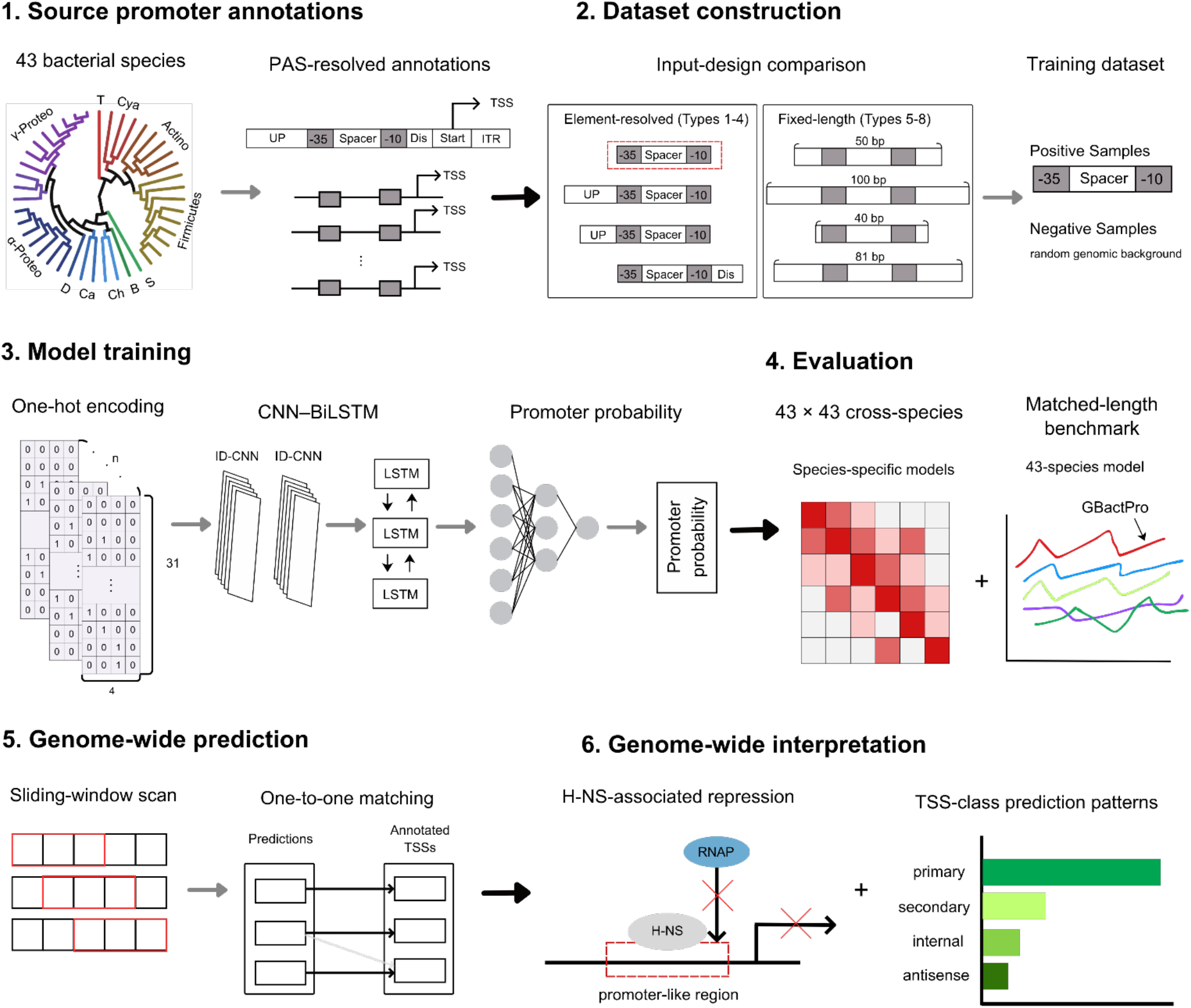
Data-centric GBactPro workflow. The workflow comprises six steps. Step 1 obtains PAS-resolved promoter annotations from 43 bacterial species. Step 2 compares four element-resolved and four fixed-length input designs, identifies Type 1 as the optimal representation, and uses this design to construct the positive (*pos_sigma70_core*) and negative (*neg_background*) datasets. Step 3 one-hot encodes each sequence as a 4 × 31 matrix and processes it through two convolutional blocks, each containing parallel one-dimensional convolutional layers with kernel sizes of 3 and 5, followed by a bidirectional LSTM layer and fully connected layers to generate promoter-prediction scores. Step 4 evaluates cross-species generalization using species-specific models across all 43 × 43 training–testing combinations and benchmarks GBactPro trained on data from all 43 species against four existing promoter predictors. Step 5 applies GBactPro across the genome using a sliding-window scan and matches predicted windows one-to-one with reference promoters for site-level evaluation. Step 6 interprets genome-wide predictions through analyses of H-NS association and TSS-class differences.

## Materials and Methods

### Source promoter annotations and species filtering

We obtained element-resolved promoter annotations for 49 bacterial species from Kuo et al. (9). Experimentally mapped TSS datasets from multiple sources were curated to identify primary TSSs and analyzed using PAS to resolve promoter architecture consistently across species. Six species were excluded because their reduced model performance in Kuo et al. raised concerns about the reliability of the underlying TSS mapping data, leaving 43 species for subsequent analyses (Supplementary Table S1).

### Input-sequence designs

To examine the influence of input-sequence definition on cross-species prediction, we evaluated eight input designs (Types 1–8; Table 1). Types 1–4 were PAS-anchored, element-resolved designs. Type 1 comprised the contiguous −35/spacer/−10 core, whereas Types 2 and 3 extended this core upstream by 50 and 20 bp, respectively, toward the UP-element region. Type 4 additionally incorporated the discriminator downstream of the −10 element. Types 5–8 used fixed-length promoter windows: Types 5 and 6 were 50- and 100-bp PAS-anchored extensions of the Type 1 core on both sides, whereas Types 7 and 8 were TSS-anchored windows. Type 7 comprised the 40 bp upstream of the TSS, matching the input used by Promotech (11); Type 8 spanned TSS −60 to +21 (81 bp), matching the input lengths used by iPro-MP (12), Prompt (15), and iPro-WAEL (14). Thus, the eight designs distinguish element-resolved promoter definitions from fixed-length windows, without treating all fixed-length designs as TSS-anchored.

**Table 1.** Definitions of the eight input-sequence types.

| Type | Input design | Length (bp) | Sequence composition |
| --- | --- | --- | --- |
| 1 | Element-resolved | 27–31 | $-35 + \text{spacer} + -10$ |
| 2 | Element-resolved | 77–81 | UP (50 bp) + $-35 + \text{spacer} + -10$ |
| 3 | Element-resolved | 47–51 | UP (20 bp) + $-35 + \text{spacer} + -10$ |
| 4 | Element-resolved | 27–44 | $-35 + \text{spacer} + -10 + \text{discriminator (DIS, 0–13 bp)}$ |
| 5 | Fixed-length | 50 | $-35 + \text{spacer} + -10$, extended on both sides |
| 6 | Fixed-length | 100 | $-35 + \text{spacer} + -10$, extended on both sides |
| 7 | Fixed-length | 40 | 40 bp upstream of TSS |
| 8 | Fixed-length | 81 | TSS $-60$ to $+21$ |

### Positive and negative sample construction

Positive samples were generated from experimentally mapped TSSs according to the eight input designs. Negative samples were randomly sampled from genomic regions at least 200 bp away from annotated promoters and matched to the corresponding positive-sequence length distributions. The default Type 1 positive dataset, *pos_sigma70_core*, comprised contiguous −35/spacer/−10 sequences of 27–31 bp; the corresponding negative dataset was designated *neg_background*. Within each species, positive sequences were divided into 75% training and 25% held-out testing sets, and negative samples were then selected separately for each partition. The training sets used a 1:10 positive-to-negative ratio to provide broader background coverage, whereas the test sets used a balanced 1:1 ratio for consistent evaluation across species (Supplementary Table S1).

### Model architecture and training

Sequences were one-hot encoded as four nucleotide channels. Type 1 inputs, ranging from 27 to 31 bp, were zero-padded at the 5′ end to 31 bp, maintaining alignment of the −10 element at the 3′ end and yielding 4 × 31 input matrices. Preliminary random-forest models were sensitive to changes in the relative positions of the −35 and −10 elements caused by variable spacer length and were therefore not retained. We adopted a CNN–BiLSTM architecture adapted from DeePromClass (28). The network contained two convolutional blocks, each with parallel one-dimensional convolutional layers with kernel sizes of 3 and 5. The first and second blocks used 32 and 64 filters, respectively, followed by batch normalization, max pooling, and dropout (0.5). A bidirectional long short-term memory (BiLSTM) layer with 1,024 units was followed by a dense layer with 1,024 units and a sigmoid output for promoter prediction. Models were trained with the Adam optimizer (learning rate 0.001), binary cross-entropy loss, and batch size 32. Complete architectural specifications are provided in Supplementary Table S2.

### Cross-species evaluation and model benchmarking

Prediction performance was evaluated using the area under the receiver operating characteristic curve (AUROC) and the area under the precision–recall curve (AUPRC). For the 43 × 43 species-specific transfer evaluation, each model was trained on the training partition of one bacterial species and evaluated on the held-out test partitions of all 43 species. The diagonal of each matrix represents within-species evaluation, whereas off-diagonal entries represent transfer between species. Performance was summarized by row-average values for training species and column-average values for testing species.

For benchmarking against existing predictors, GBactPro was also trained jointly on the pooled training sets of all 43 species and evaluated on their held-out test sets. Alongside the default Type 1 (27–31 bp) input, separate jointly trained GBactPro models used the matched-length Type 8 (81 bp) input for comparison with iPro-MP (12), Prompt (15), and iPro-WAEL (14), and the Type 7 (40 bp) input for comparison with Promotech (11). Models trained exclusively on *E. coli* were evaluated as additional controls to distinguish input-design effects from the benefits of joint multi-species training. All tools were evaluated on the same held-out test sequences, with mean performance across the 43 testing species and separate *E. coli* performance reported. Model architectures, training scopes, and input designs are summarized in Supplementary Table S3.

### Genome-wide prediction

For genome-wide prediction, GBactPro was retrained on the pooled 43-species training dataset using a 29-bp configuration adapted from the Type 1 (−35/spacer/−10) input design. Only promoters with spacer lengths of 17 bp or shorter were retained, yielding 27–29-bp sequences that were zero-padded at the 5′ end to 29 bp while maintaining alignment of the −10 element at the 3′ end. The 29-bp window was selected to accommodate the most common spacer length of 17 bp without requiring the 31-bp input used for the complete Type 1 dataset. Both strands of the approximately 4.6-Mb *E. coli* genome were scanned using 29-bp windows at a one-nucleotide stride, generating approximately 9.2 million strand-specific windows, each assigned a promoter-prediction score. For evaluation, a 34-bp reference promoter region was defined for each TSS as [TSS − 39, TSS − 6] on the positive strand and [TSS + 6, TSS + 39] on the negative strand. These regions encompass the expected −35/spacer/−10 core, allowing spacer lengths of 15–19 bp and a distance of 6–9 bp between the 3′ end of the −10 element and the TSS (Supplementary Fig. S1).

To evaluate promoter-site detection, predicted windows were matched to reference promoter regions using a score-ordered, one-to-one greedy procedure (Supplementary Fig. S2). At each prediction-score threshold, windows meeting the threshold were processed in descending order of GBactPro score. Under the stringent 0-bp tolerance criterion, a predicted window was eligible for matching only if it was fully contained within a reference promoter region on the same strand. Each predicted window and reference TSS could be matched at most once. When multiple windows corresponded to the same reference promoter, the highest-scoring window was matched first, whereas lower-scoring windows could match only remaining unmatched reference promoters. If a window corresponded to multiple unmatched reference promoter regions, it was assigned to the closest reference TSS. Matched predictions were classified as true positives (TPs), predictions without an assigned reference promoter as false positives (FPs), and unmatched reference TSSs as false negatives (FNs). This procedure prevented multiple overlapping predictions from being counted as separate true positives for the same promoter.

Genome-wide performance was assessed using AUROC, AUPRC, false positives per kilobase, and precision at ranked score cutoffs. Because promoter-associated windows are greatly outnumbered by non-promoter windows in genome-wide scanning, AUPRC was reported alongside AUROC to better characterize performance under class imbalance (29). ROC and precision–recall curves were generated across prediction-score thresholds using the one-to-one matching procedure and integrated using NumPy trapezoidal integration. For site-level evaluation, the threshold maximizing the F1 score was selected under each matching criterion. A random baseline was generated from 1,000 simulations using randomly assigned prediction scores and the same matching procedure. To account for predictions near annotated promoter regions, a relaxed 100-bp tolerance was also evaluated by extending each reference promoter region by 100 bp in both directions while retaining the one-to-one matching procedure.

### H-NS binding and knockout datasets

To investigate the association between GBactPro predictions and H-NS-associated transcriptional repression, we analyzed published H-NS binding and knockout datasets from *E. coli*. For the binding analysis, we examined the overlap between genome-wide GBactPro predictions and 547 H-NS binding regions identified by ChIP-seq (NCBI GSE24991) (30). A contingency-table χ^2^ test was used to assess the association between prediction outcome (TP versus FP) and H-NS overlap. Among FPs, prediction scores were compared between H-NS-overlapping and non-overlapping predictions using a Mann–Whitney U test. Both analyses used predictions at the genome-wide best-F1 threshold under the 0-bp tolerance criterion.

For the knockout analysis, we used an independent published *Δhns* dataset containing 668 high-confidence TSSs colocalized with σ70 ChIP-seq peaks (31). For each TSS, a reference promoter region was assigned using the definition established for genome-wide evaluation. TSS strength was defined in the original study by RNA-seq read density at the TSS. Promoters were classified as derepressed when their TSS strength was higher in the *Δhns* strain than in the wild-type strain. Derepressed promoters were evaluated for overlap with GBactPro predictions under the 0- and 100-bp tolerance criteria.

### Classification and curation of *E. coli* transcription start sites

To evaluate GBactPro predictions across different transcriptional contexts, we used the *E. coli* TSS annotations from Thomason et al. (32), comprising four classes: (i) primary TSSs, the main transcription initiation sites of genes or operons; (ii) secondary TSSs, alternative initiation sites with lower expression than their corresponding primary TSSs; (iii) internal TSSs, located within annotated genes on the sense strand; and (iv) antisense TSSs, located within or near annotated genes on the opposite strand.

Only TSSs labeled as both “enriched” and “detected” were retained. Internal and antisense TSSs were additionally required to have *detCount* ≥ 2, indicating detection in at least two experimental conditions. After filtering, the four classes contained 2,672 primary, 1,589 secondary, 4,246 internal, and 4,763 antisense TSSs. Because some TSSs were assigned to multiple classes, overlapping class assignments were resolved using the priority order primary > secondary > internal > antisense, resulting in 11,834 uniquely classified TSSs.

## Results

### Element-resolved promoter definition improves cross-species prediction

To assess the effect of promoter-sequence definition on cross-species prediction, we trained GBactPro models on *E. coli* data using each of the eight input designs and evaluated them on held-out test sets from all 43 bacterial species (Table 2; Supplementary Fig. S3). All four element-resolved designs (Types 1–4) achieved higher mean AUROC, AUPRC, and accuracy than the four fixed-length designs (Types 5–8). The element-resolved designs achieved mean AUROC values of 0.9602–0.9859, compared with 0.6984–0.9064 for the fixed-length designs. Even the lowest-performing element-resolved design (Type 3, UP (20 bp)/−35/spacer/−10; AUROC = 0.9602) exceeded the best fixed-length design (Type 7, 40 bp upstream of the TSS; AUROC = 0.9064).

**Table 2.** Performance of the eight input-sequence types.

| Type | Sequence composition | Length (bp) | Mean AUROC | Mean AUPRC | Mean Accuracy |
| --- | --- | --- | --- | --- | --- |
| 1 | –35 + spacer + –10 | 27–31 | 0.9859 | 0.9853 | 0.9477 |
| 2 | UP (50 bp) + –35 + spacer + –10 | 77–81 | 0.9633 | 0.9647 | 0.9115 |
| 3 | UP (20 bp) + –35 + spacer + –10 | 47–51 | 0.9602 | 0.9606 | 0.9087 |
| 4 | –35 + spacer + –10 + DIS | 27–44 | 0.9738 | 0.9736 | 0.9271 |
| 5 | Type 1 extended on both sides | 50 | 0.9056 | 0.9048 | 0.8427 |
| 6 | Type 1 extended on both sides | 100 | 0.6984 | 0.6959 | 0.6814 |
| 7 | 40 bp upstream of TSS | 40 | 0.9064 | 0.9101 | 0.8470 |
| 8 | TSS -60 to +21 | 81 | 0.8398 | 0.8284 | 0.7761 |

Among the element-resolved designs, Type 1 (−35/spacer/−10) performed best, followed by Types 4, 2, and 3, with consistent rankings across all three evaluation metrics. Type 1 achieved mean AUROC, AUPRC, and accuracy values of 0.9859, 0.9853, and 0.9477, respectively, and was selected as the default GBactPro input for subsequent analyses.

Among the fixed-length designs, Type 7 (40 bp; AUROC = 0.9064) and Type 5 (50 bp; AUROC = 0.9056) showed similar performance, followed by Type 8 (81 bp; AUROC = 0.8398) and Type 6 (100 bp; AUROC = 0.6984). This ranking was consistent across AUROC, AUPRC, and accuracy. The lower performance of the longer fixed-length designs suggests that incorporating additional sequence beyond the core promoter does not necessarily improve cross-species prediction.

### GBactPro generalizes across 43 bacterial species

To evaluate cross-species transfer, we trained 43 species-specific GBactPro models and evaluated each model on the held-out test sets of all 43 bacterial species, generating a 43 × 43 performance matrix (Fig. 2A; Supplementary Fig. S4A). Pairwise AUROC values ranged from approximately 0.70 to greater than 0.99, with row-average and column-average AUROC values ranging from 0.882 to 0.993 and from 0.929 to 0.994, respectively (Supplementary Table S4). Although performance varied among training–testing species pairs, most models maintained high predictive performance across species. Model performance was also associated with the number of available promoter annotations in the training species. Row-average AUROC and AUPRC increased rapidly with promoter count for smaller datasets and approached saturation as dataset size increased (Fig. 2B; Supplementary Fig. S4B). The fitted curves explained substantial variation in row-average performance (R^2^ = 0.815 for AUROC and 0.802 for AUPRC), indicating a positive association between promoter dataset size and cross-species prediction performance.

**Figure 2.**
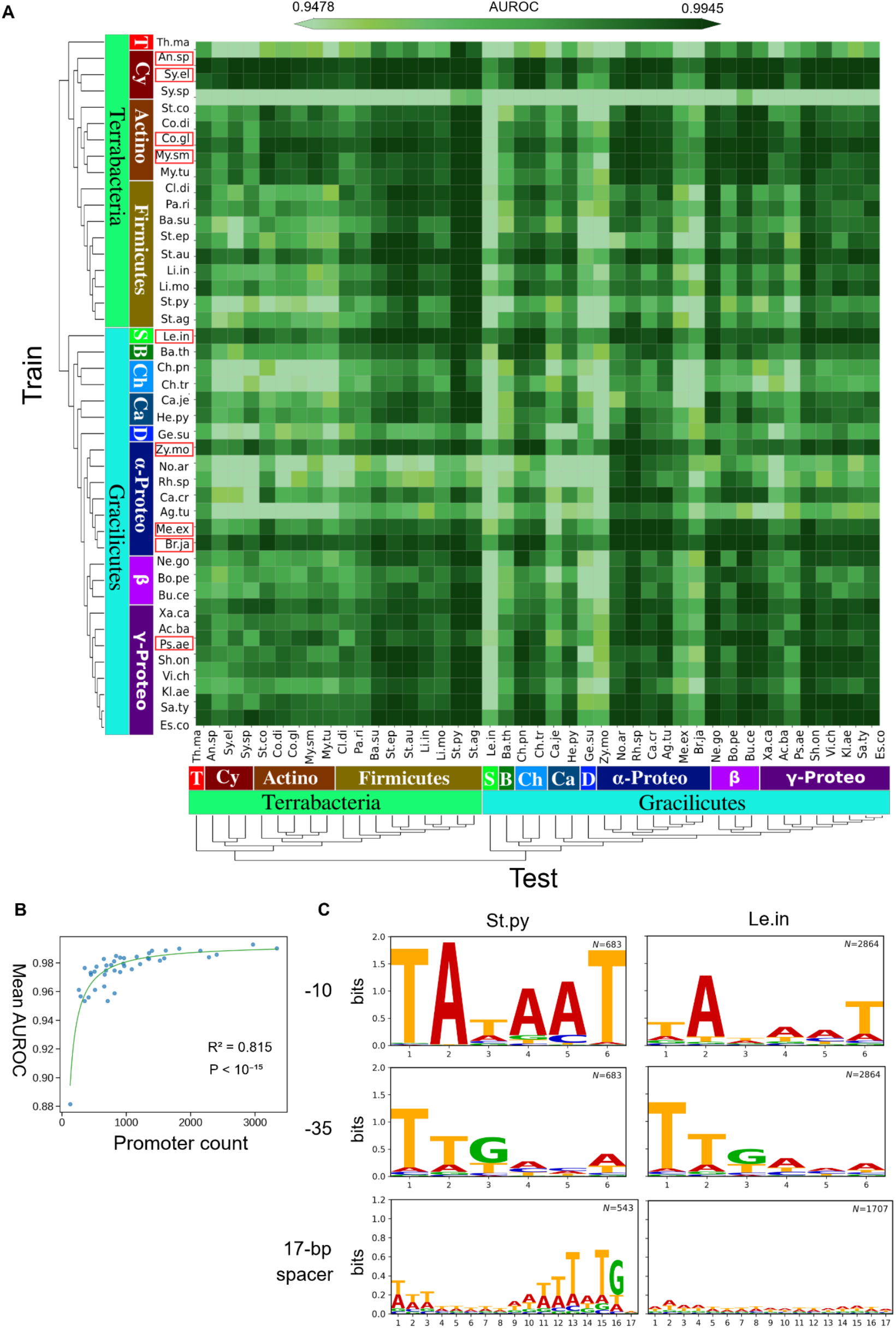
Cross-species generalization across 43 bacterial species. (A) 43 × 43 AUROC heatmap of species-specific models evaluated across all 43 species. Rows represent training species and columns represent testing species, with species ordered by lineage. Species with >2,000 total annotated promoters (before the training/test split) are marked with red boxes. (B) Relationship between training promoter count and the mean cross-species performance of each species-specific model across all 43 testing species, corresponding to the row-average AUROC in A. Each point represents one species-specific model, and the green curve indicates the saturating fit. (C) Sequence logos for *Streptococcus pyogenes* and *Leptospira interrogans*, the species with the strongest and weakest recognition across the 43 species-specific models, respectively, based on the highest and lowest column-average AUROC in A. The 17-bp spacer represented approximately 79.5% (543/683) and 60% (1707/2864) of promoters in *S. pyogenes* and *L. interrogans*, respectively.

To investigate differences in the recognition of testing species, we compared promoter-sequence conservation between *Streptococcus pyogenes* and *Leptospira interrogans*, which showed the highest and lowest column-average AUROC values, respectively (Supplementary Table S4). Sequence-logo analysis (Fig. 2C) revealed stronger conservation of the −10 element and more distinct spacer-associated motifs in *S. pyogenes*, including an upstream AT-rich tract, a central AT-rich tract, and a TG motif positioned one base pair upstream of the −10 element. In contrast, *L. interrogans* showed weaker conservation of the −10 element and less distinct spacer-associated motifs, whereas differences in the −35 element were less pronounced. These observations suggest that conservation of core promoter sequence features may contribute to differences in cross-species recognition performance.

### GBactPro outperforms existing predictors under matched-length benchmarks

A GBactPro model trained jointly on pooled data from all 43 species outperformed the corresponding species-specific models in 41 of 43 testing species by AUROC and 37 of 43 by AUPRC (Supplementary Fig. S5). Using the default Type 1 (−35/spacer/−10) input, the jointly trained model achieved a mean AUROC of 0.9943 across the held-out test sets of all 43 species (Table 3). To compare GBactPro with existing predictors under matched input lengths, we evaluated separately trained GBactPro models using Type 8 (81 bp) and Type 7 (40 bp) inputs. At 81 bp, GBactPro achieved a mean AUROC of 0.9431, exceeding iPro-MP (0.8416), Prompt (0.7784), and iPro-WAEL (0.6293). At 40 bp, GBactPro achieved a mean AUROC of 0.9426, compared with 0.8757 for Promotech. GBactPro also achieved higher mean AUPRC values than the corresponding predictors under both matched-length settings, with the same performance advantage observed on the *E. coli* test set (Table 3; Supplementary Fig. S7).

**Table 3.** Cross-species performance and matched-length benchmark comparison using the same held-out test sequences.

| Tool | Input length | Training data | Mean |  | *E. coli* |  |
| --- | --- | --- | --- | --- | --- | --- |
|  |  |  | AUROC | AUPRC | AUROC | AUPRC |
| GBactPro | 27–31 bp (Type 1) | 43 species | 0.9943 | 0.9931 | 0.9968 | 0.9964 |
| GBactPro | 27–31 bp (Type 1) | *E. coli* | 0.9859 | 0.9853 | 0.9926 | 0.9921 |
| GBactPro | 81 bp (Type 8) | 43 species | 0.9431 | 0.9452 | 0.9557 | 0.9576 |
| GBactPro | 81 bp (Type 8) | *E. coli* | 0.8398 | 0.8284 | 0.8688 | 0.8405 |
| iPro-MP | 81 bp | *E. coli* | 0.8416 | 0.8472 | 0.9059 | 0.9170 |
| Prompt | 81 bp | *E. coli* | 0.7784 | 0.7645 | 0.7850 | 0.7568 |
| iPro-WAEL | 81 bp | *E. coli* | 0.6293 | 0.6202 | 0.6802 | 0.6945 |
| GBactPro | 40 bp (Type 7) | 43 species | 0.9426 | 0.9495 | 0.9563 | 0.9605 |
| GBactPro | 40 bp (Type 7) | *E. coli* | 0.9064 | 0.9101 | 0.9290 | 0.9246 |
| Promotech | 40 bp | 9 species | 0.8757 | 0.8750 | 0.9071 | 0.9037 |

To assess the influence of training-data scope, we further evaluated GBactPro models trained exclusively on *E. coli* data. The element-resolved Type 1 model achieved a mean AUROC of 0.9859 across the 43 testing species, compared with 0.9064 and 0.8398 for the fixed-length Type 7 and Type 8 models, respectively. Under the 81-bp setting, the *E. coli*-trained GBactPro model achieved a mean AUROC of 0.8398, exceeding Prompt (0.7784) and iPro-WAEL (0.6293), but slightly below iPro-MP (0.8416). Under the 40-bp setting, the *E. coli*-trained GBactPro model achieved a mean AUROC of 0.9064, exceeding Promotech (0.8757), which was trained on data from nine species. Together, these results support the benefits of PAS-resolved promoter-core inputs and indicate that joint multi-species training can further improve prediction performance.

### Genome-wide promoter-site prediction in *E. coli*

We next applied GBactPro to genome-wide promoter-site prediction in *E. coli*, using overlapping 29-bp windows to identify candidate promoter sites. *E. coli* was selected because its extensive TSS annotations and regulatory datasets support site-level evaluation and subsequent biological interpretation. Under the stringent 0-bp tolerance criterion with one-to-one matching, GBactPro achieved a genome-wide AUROC of 0.979 and an AUPRC of 0.160 (Fig. 3A), with the latter approximately 30-fold higher than the random baseline of 0.0056. At the best-F1 prediction-score threshold of 0.9315 under the 0-bp tolerance criterion, GBactPro produced 2.556 false positives per kilobase. When predictions were ranked by GBactPro score, precision reached approximately 0.80 among the top 25 predictions and 0.72 among the top 100. These results indicate that GBactPro maintained strong genome-wide discrimination, with annotated promoters enriched among high-scoring predictions despite the remaining false-positive burden.

**Figure 3.**
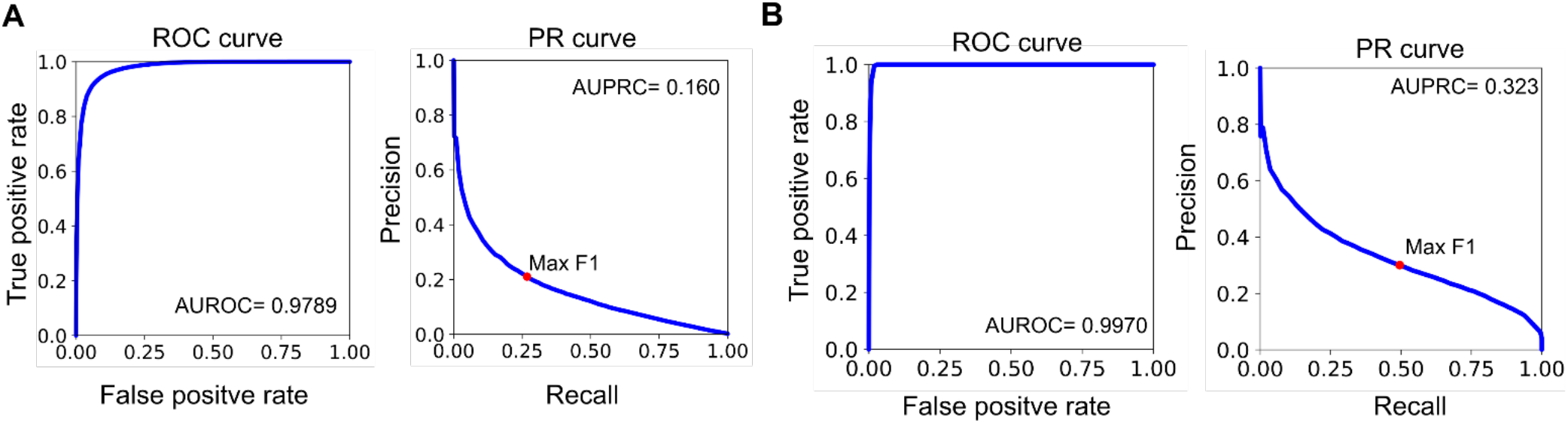
Promoter detection across the *E. coli* genome under different tolerance settings. (A) ROC and precision–recall curves under the stringent 0-bp tolerance criterion, requiring predicted windows to be fully contained within a same-strand 34-bp reference promoter region and assigned using one-to-one matching. (B) Corresponding curves under the relaxed 100-bp tolerance criterion. Red dots indicate the threshold maximizing the F1 score.

To assess whether some unmatched predictions were located near annotated promoters, we repeated the evaluation using a relaxed 100-bp tolerance while retaining the one-to-one matching procedure. Under this criterion, AUROC increased to 0.997 and AUPRC to 0.323 (Fig. 3B). Sequence-logo analysis further showed that promoters associated with unmatched reference TSSs exhibited weaker conservation of the −10 element and spacer-associated motifs than those associated with matched TSSs (Supplementary Fig. S6). This difference suggests that reduced conservation of core promoter features may contribute to missed predictions. A direct genome-wide benchmark against existing predictors was not performed because their required 40- and 81-bp inputs could not satisfy the stringent full-containment criterion within the 34-bp reference promoter regions without modifying the evaluation framework.

### Association between GBactPro predictions and H-NS binding

To investigate whether unmatched GBactPro predictions were associated with transcriptional repression, we examined their overlap with H-NS binding regions in *E. coli*. H-NS is a nucleoid-associated protein that preferentially binds AT-rich DNA and represses transcription from many bacterial promoters (30,33). At the genome-wide best-F1 threshold under the stringent 0-bp tolerance criterion, FPs showed significantly greater overlap with H-NS binding regions than TPs (24.4% versus 18.8%; χ^2^ test, P < 0.001; Fig. 4A). Among FPs, H-NS-overlapping predictions also received higher scores than non-overlapping predictions (Mann–Whitney U test, P = 0.027). At a representative genomic locus, several high-scoring predicted windows overlapped an H-NS binding region without a corresponding annotated reference TSS (Fig. 4B).

**Figure 4.**
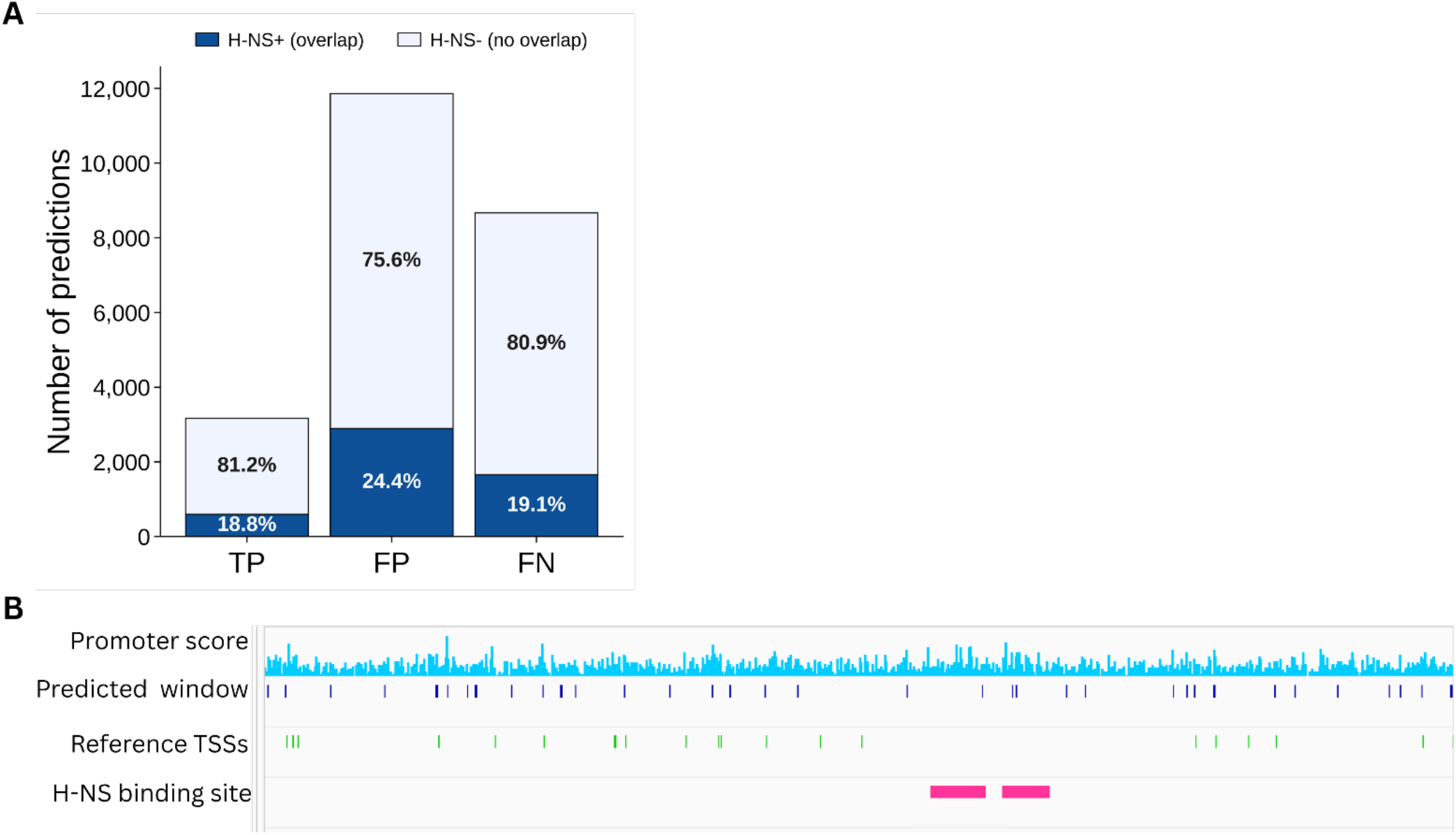
Association between GBactPro predictions and H-NS binding. (A) Stacked counts of true positives, false positives, and false negatives under the stringent 0-bp matching criterion (F1 = 0.236; prediction-score threshold = 0.9315), partitioned by H-NS overlap. (B) Representative genome-locus view showing the prediction-score track on the plus strand (cyan, top row), predicted windows (blue, second row), reference TSSs from all four classes (green, third row), and annotated H-NS binding sites (pink, bottom row), including HNS_site_829.

We further examined an independent H-NS-knockout (*Δhns*) dataset to assess the recovery of promoters associated with H-NS repression. Among 371 derepressed promoters, 63.3% overlapped at least one GBactPro prediction under the 0-bp tolerance criterion, increasing to 90.3% under the relaxed 100-bp criterion. Together, these findings indicate an association between unmatched promoter predictions and H-NS occupancy, consistent with the possibility that some apparent false positives represent promoter-like regions whose transcriptional activity is suppressed by H-NS.

### Prediction patterns differ across TSS classes

Because GBactPro was trained on promoters derived from primary TSSs, we examined its ability to identify promoters associated with four TSS classes: primary, secondary, internal, and antisense. Under the 100-bp tolerance criterion, the distributions of TPs, FPs, and FNs differed across these classes (Fig. 5A). Primary TSSs showed the highest overall precision (0.190), followed by antisense (0.144), internal (0.123), and secondary TSSs (0.078). Despite their lower precision, internal and antisense TSSs showed higher recall than primary TSSs, indicating that a greater proportion of their reference promoters was recovered, although with a higher false-positive burden.

**Figure 5.**
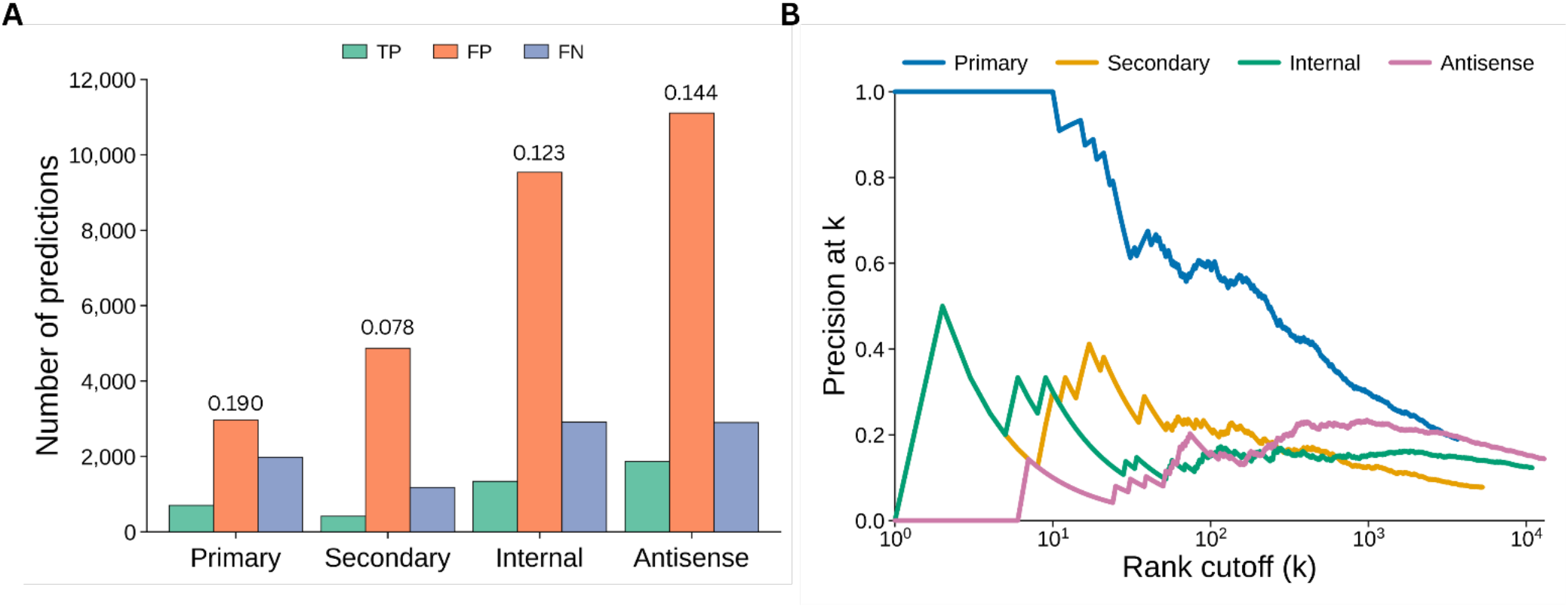
Genome-wide prediction patterns across four TSS classes. (A) TP/FP/FN counts for primary, secondary, internal, and antisense TSSs under the 100-bp tolerance criterion. Values above the bars indicate overall precision (TP/[TP + FP]). (B) Precision-at-rank curves for the four TSS classes. Colored curves show precision among the top-k predictions ranked by prediction score for each TSS class, using one-to-one greedy matching under the 100-bp tolerance criterion.

Precision-at-rank analysis further revealed distinct prediction patterns across TSS classes (Fig. 5B). Primary TSSs showed the strongest ranking performance, reaching a precision of 0.80 among the top 20 predictions. Although secondary TSSs had the lowest overall precision, their top 20–100 predictions showed higher precision than those associated with internal and antisense TSSs. For internal TSSs, precision generally declined toward the overall class-level precision as lower-ranked predictions were included. In contrast, antisense TSSs showed low precision among the highest-ranked predictions, with precision increasing as additional lower-ranked predictions were included. Together, these findings indicate that GBactPro can recover promoters associated with multiple TSS classes, but its ability to prioritize high-confidence predictions differs substantially among transcriptional contexts.

## Discussion

Our results demonstrate that promoter-sequence definition is an important determinant of bacterial promoter prediction performance. Building on the PAS-derived promoter annotations of Kuo et al. (9), GBactPro uses the contiguous −35/spacer/−10 core as its training input, rather than treating the promoter as a fixed-length window relative to the TSS. Among the eight input designs evaluated, the PAS-resolved core achieved the highest mean prediction performance across 43 bacterial species. This finding extends the biological characterization of promoter architecture into a sequence-based predictive framework that does not require prior TSS information at inference time. Compared with conventional fixed-window definitions, the element-resolved approach limits the input to core promoter elements while reducing surrounding sequence that may contribute less to promoter recognition. Although differences in input length and sequence composition may also contribute to the observed performance gains, these results demonstrate the importance of biologically informed training-sequence definitions.

The benefits of element-resolved promoter training extended across phylogenetically diverse bacterial species. The 43 × 43 evaluation revealed broad cross-species transfer, with performance associated with both training promoter count and the conservation of core promoter sequence features. Joint training on data from all 43 species further improved prediction relative to species-specific models for most testing species, achieving a mean AUROC of 0.9943 on held-out sequences. Under matched-length benchmarks, the jointly trained GBactPro models also outperformed four established predictors, including the DNABERT-based iPro-MP (11,12,14,15). The benchmark comparisons also differed in training-data scope. Under *E. coli*-only training, GBactPro performed comparably to iPro-MP at 81 bp, while retaining higher AUROC than Prompt, iPro-WAEL, and Promotech under their respective matched-length settings. Together, these findings indicate that both promoter-sequence definition and multi-species training contribute to predictive performance, while demonstrating that a CNN–BiLSTM architecture can effectively learn transferable promoter features without large-scale sequence pretraining.

Beyond sequence-level classification, GBactPro was applied to genome-wide promoter-site prediction in *E. coli* without requiring prior TSS annotations during prediction. Using a one-to-one matching procedure to prevent redundant true-positive assignments, GBactPro achieved an AUROC of 0.979 and an AUPRC of 0.160 under stringent positional matching. These values increased to 0.997 and 0.323, respectively, under a relaxed 100-bp tolerance. Although the relaxed criterion accounts for predictions near annotated promoter regions, it does not represent equivalent precision in promoter localization. The remaining false-positive burden also highlights the challenge of distinguishing promoter-like genomic sequences from experimentally annotated transcription initiation sites. Thus, genome-wide predictions are particularly useful for prioritizing candidate promoter regions for further investigation.

The biological interpretation of genome-wide predictions further revealed an association with H-NS-mediated transcriptional repression. Unmatched predictions were more frequently associated with H-NS binding than matched predictions, and an independent H-NS-knockout dataset showed that GBactPro recovered a substantial proportion of derepressed promoters. These observations are consistent with the possibility that some apparent false positives represent promoter-like regions whose transcriptional activity is suppressed under the reference conditions. However, H-NS binding alone does not establish functional promoter activity, and its preference for AT-rich DNA may also contribute to the observed association. Moreover, prediction performance differed across TSS classes: primary TSSs showed the strongest enrichment among high-ranking predictions, whereas secondary, internal, and antisense TSSs exhibited distinct ranking patterns. These results highlight the influence of transcriptional context on the interpretation of sequence-based promoter predictions.

Several limitations should be considered. GBactPro was trained primarily on PAS-resolved promoter cores derived from primary TSS annotations, which may favor promoters with recognizable −35/spacer/−10 architectures. Recent experimental work by Kuo et al. (34) further demonstrated distinct roles for the −35 and −10 elements in promoter formation and regulation, emphasizing the biological diversity that may not be fully represented by a single promoter-core definition. In addition, the balanced sequence-level test sets differ from the sparse promoter distribution encountered in whole-genome prediction, and genome-wide evaluation was performed only in *E. coli*. The joint 43-species evaluation also does not establish generalization to bacterial species absent from training. Finally, sequence-based predictions cannot directly account for condition-dependent transcriptional activity or regulatory protein occupancy. Within these limitations, our findings demonstrate that biologically informed promoter definitions, combined with multi-species training, provide an effective framework for cross-species bacterial promoter prediction and genome-wide identification of candidate promoter sites.

## Data Availability

GBactPro source code and documentation are publicly available at https://github.com/changlabtw/GBactPro/. Trained models, processed promoter annotations, training and testing datasets, benchmark scripts, and genome-wide prediction outputs are available through the repository and archived in Zenodo (https://doi.org/10.5281/zenodo.23718018).

## Supplementary Data

Supplementary Data are available at NAR Online.

## Author Contributions

H.Y.B.H.: Investigation, Formal analysis, Software, Writing – original draft. Y.-C.V.K.: Investigation, Formal analysis. S.T.A.K.: Writing – review & editing. H.H.D.C.: Conceptualization, Supervision, Funding acquisition, Writing – review & editing. J.-M.C.: Conceptualization, Supervision, Funding acquisition, Writing – review & editing.

## Acknowledgements

We thank the National Core Facility for Biopharmaceuticals (NCFB; NSTC 115-2740-B-492-001) and the National Center for High-performance Computing (NCHC) of the National Institutes of Applied Research (NIAR), Taiwan, for providing computational and storage resources.

