## supplemental material for "GBactPro: cross-species bacterial promoter prediction using element-resolved promoter definitions"

#### Supplementary Figures

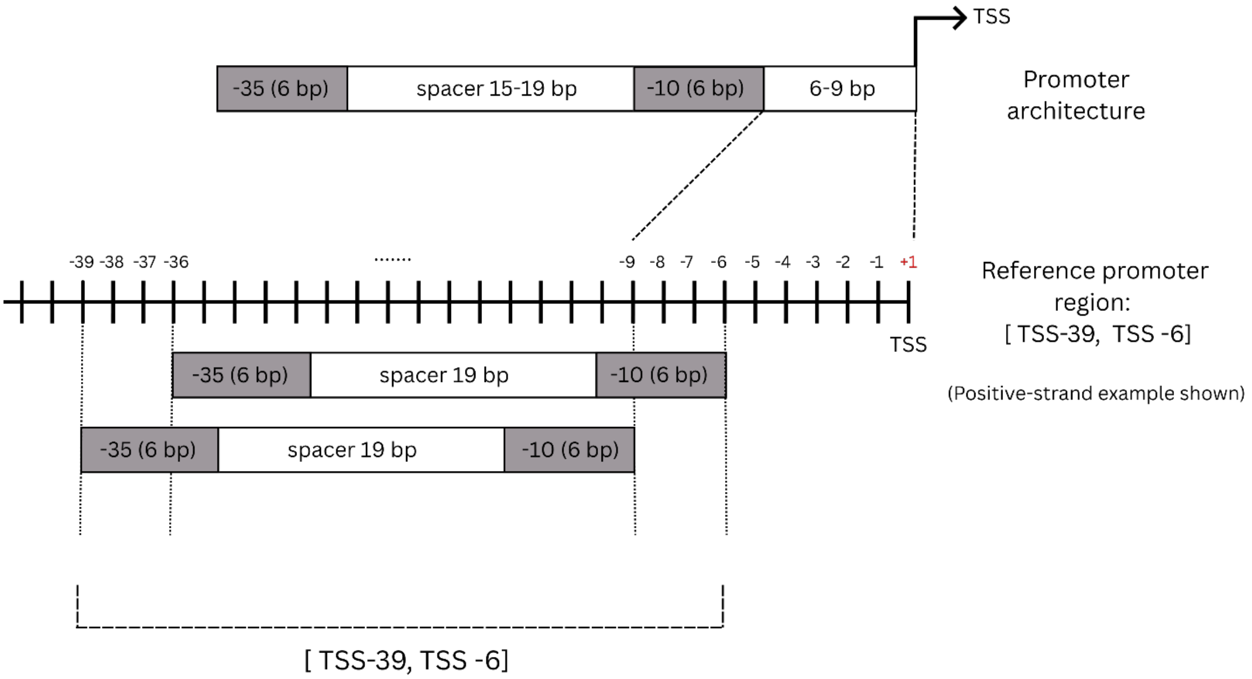


**Supplementary Figure S1. Definition of reference promoter regions for genome-wide evaluation.** Reference promoter regions were defined for each TSS as [TSS − 39, TSS − 6] on the positive strand and [TSS + 6, TSS + 39] on the negative strand, yielding a 34-bp region encompassing the expected −35/spacer/−10 core. The schematic shows the positive-strand definition. Genome-wide scanning generated 29-bp predicted windows. Under the 0-bp tolerance criterion, predicted windows were eligible for matching only when fully contained within a same-strand reference promoter region. A prediction was counted as a true positive only after assignment to an unmatched reference TSS through the one-to-one matching procedure.

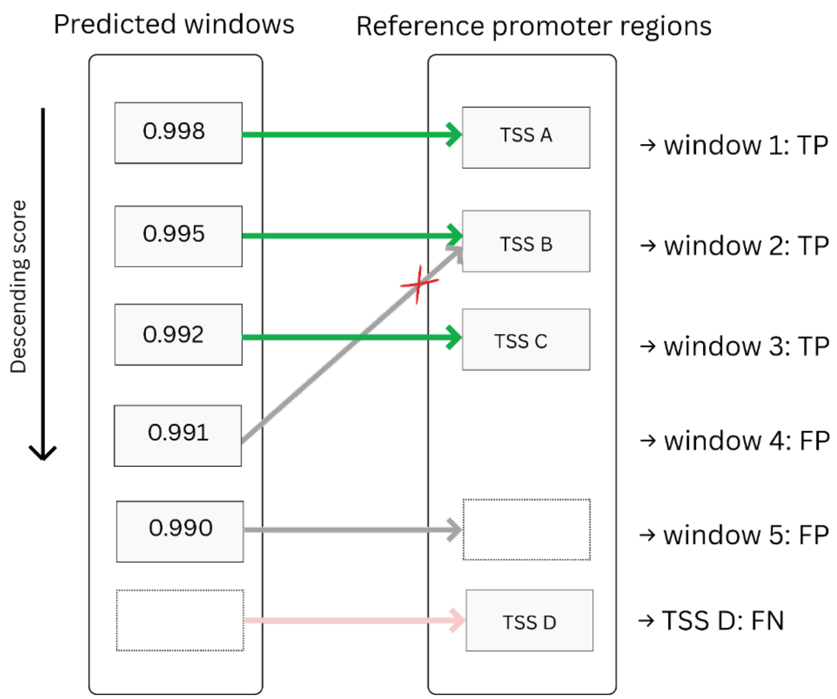


**Supplementary Figure S2. Greedy one-to-one matching of promoter sites in genome-wide evaluation.** At each prediction-score threshold, windows meeting the threshold were processed in descending order of GBactPro score. Under the stringent 0-bp criterion, only windows fully contained within a same-strand reference promoter region were eligible for matching. Each predicted window and reference TSS could be matched at most once. Lower-scoring predictions could therefore match only remaining unmatched reference TSSs. Matched predictions were classified as true positives (TPs), unmatched predictions as false positives (FPs), and unmatched reference TSSs as false negatives (FNs). Prediction scores are shown schematically.

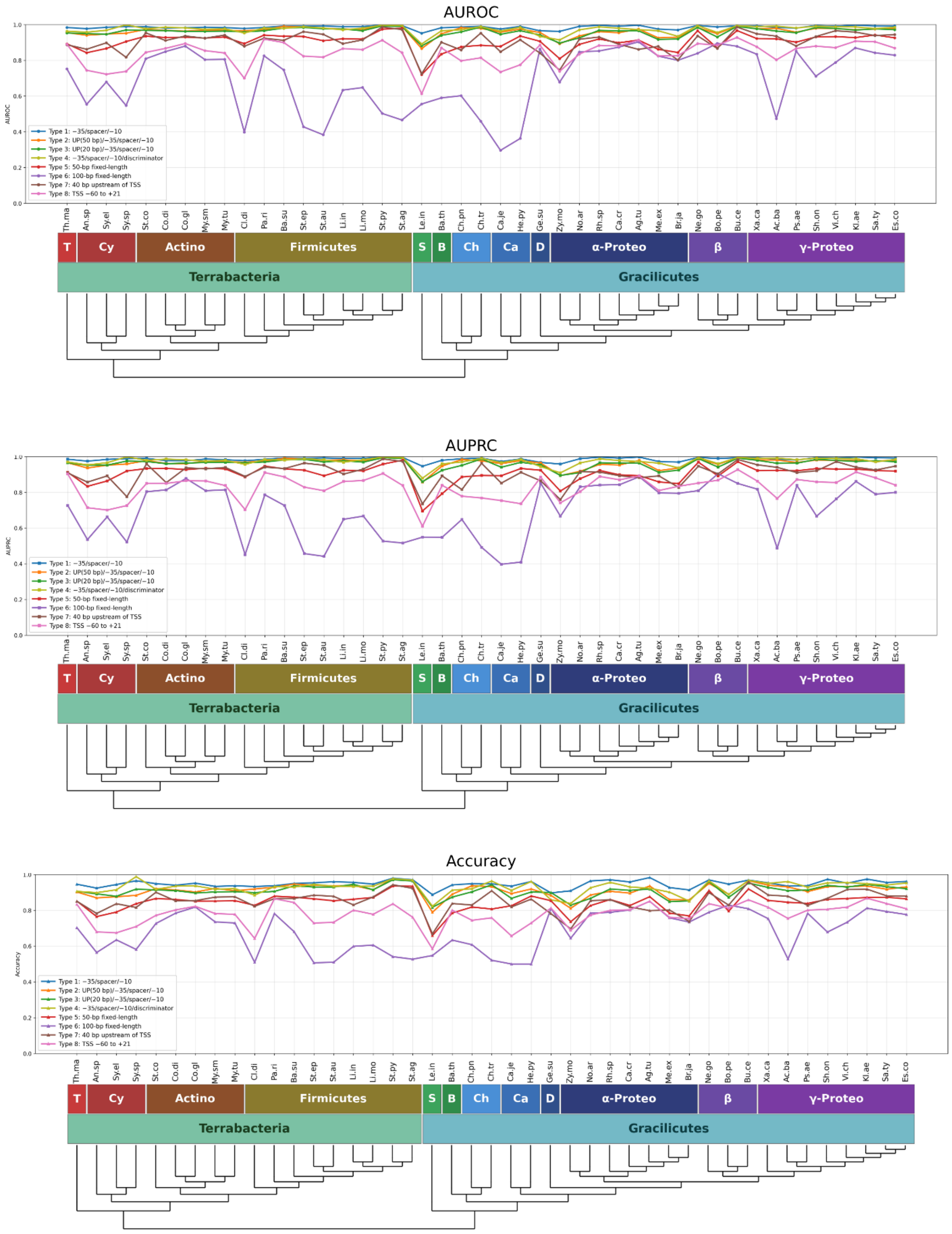


**Supplementary Figure S3. Cross-species performance of eight input-sequence designs across 43 bacterial species.** Per-species AUROC (top), AUPRC (middle), and accuracy (bottom) of CNN–BiLSTM models trained on *E. coli* using each of the eight input-sequence designs and evaluated on held-out test data from all 43 species. Species are ordered by phylogenetic relationship.

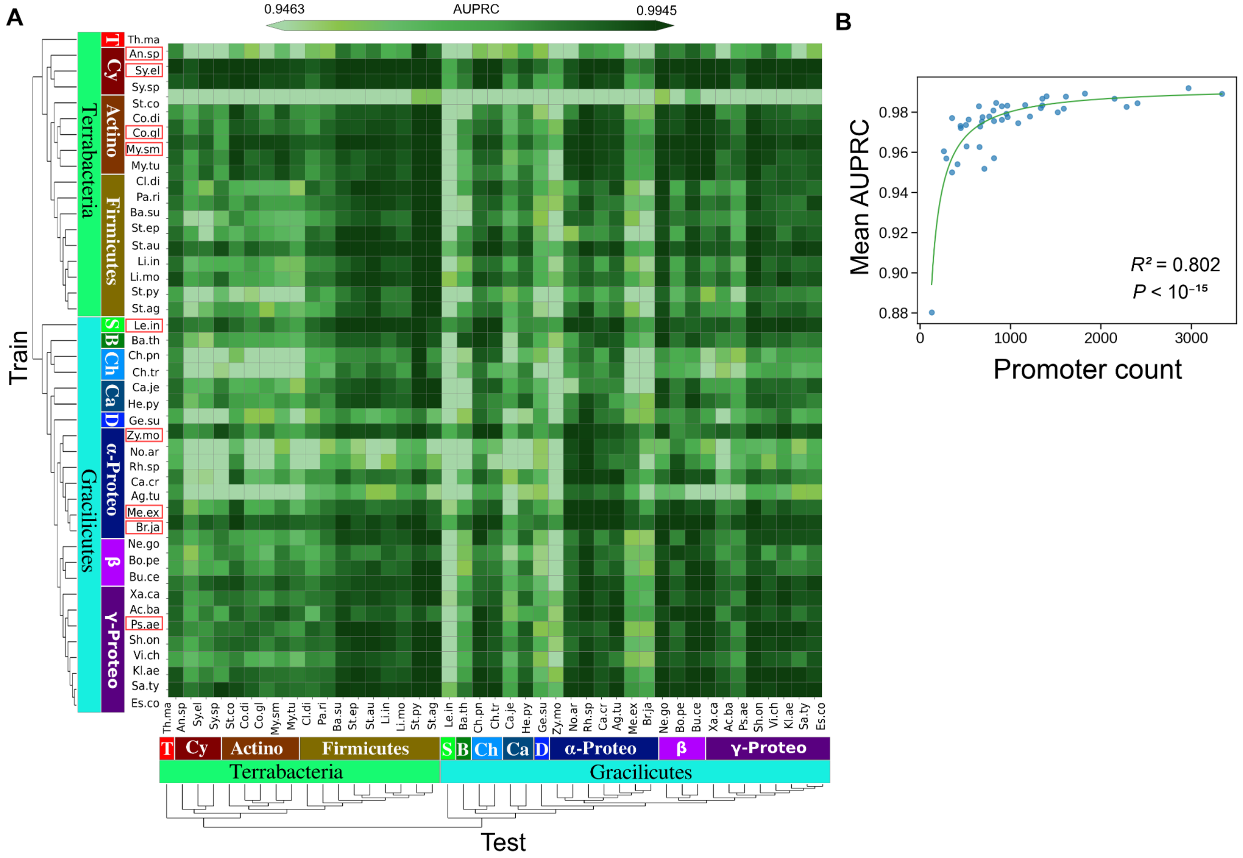


**Supplementary Figure S4. Cross-species generalization based on AUPRC across 43 bacterial species.** (A) 43 × 43 AUPRC heatmap of species-specific models evaluated across all 43 species. Rows represent training species and columns represent testing species, with species ordered by lineage. Species with >2,000 total annotated promoters (before the training/test split) are marked with red boxes. (B) Relationship between training promoter count and the mean cross-species AUPRC of each species-specific model across all 43 testing species, corresponding to the row-average AUPRC in A. Each point represents one species-specific model, and the green curve indicates the saturating fit.

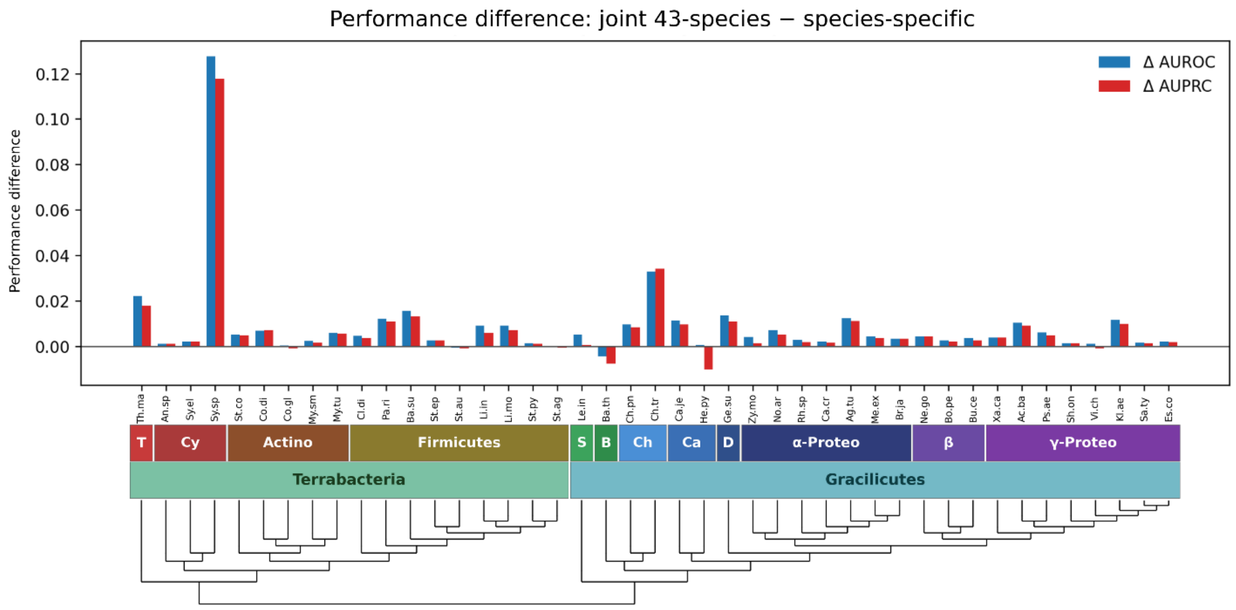


**Supplementary Figure S5. Species-level performance gains from joint 43-species training.** Per-species differences in AUROC and AUPRC between the jointly trained 43-species GBactPro model and the corresponding species-specific models. Differences were calculated as joint 43-species minus species-specific performance. Positive values indicate gains from joint training, whereas negative values indicate lower performance.

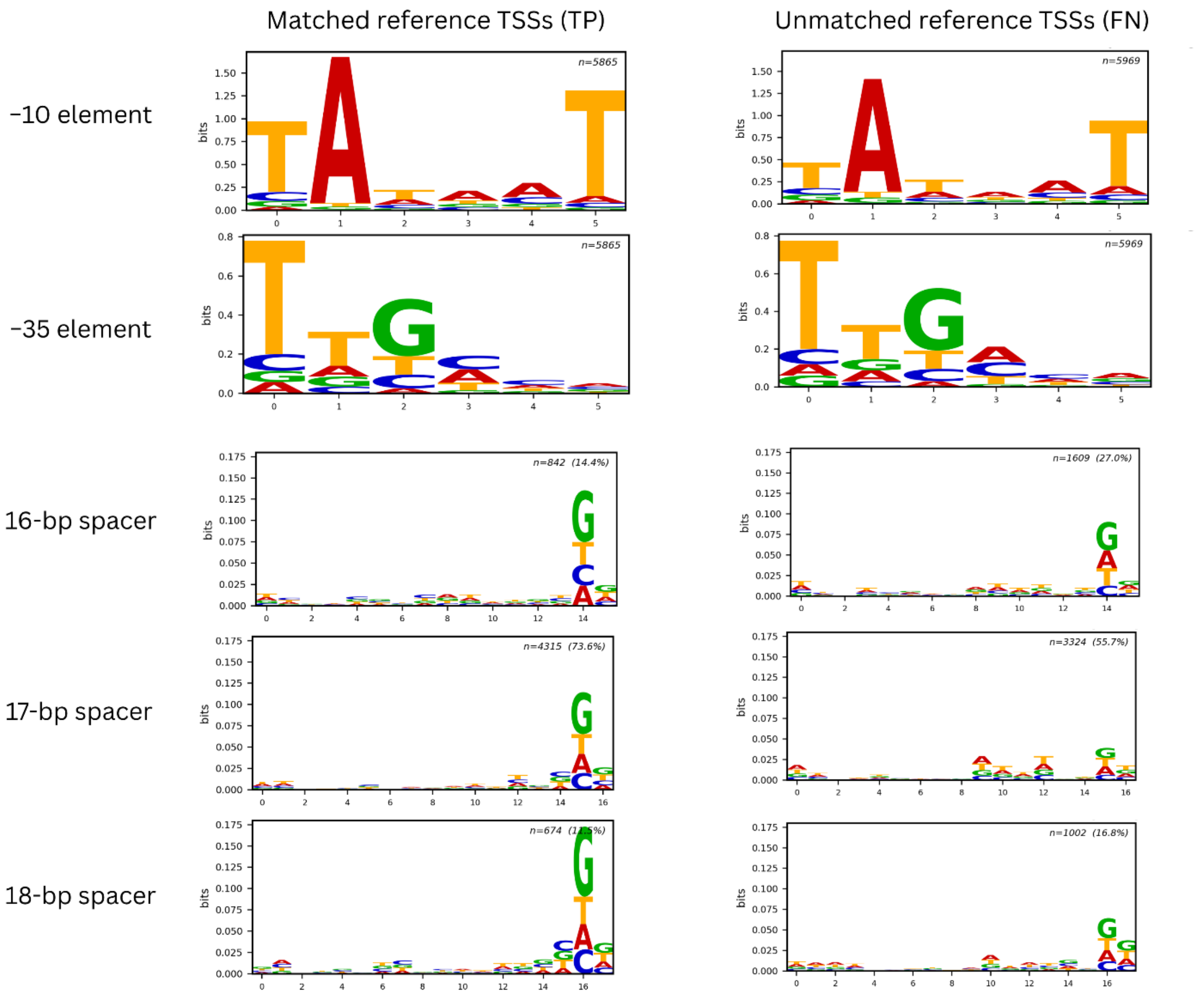


**Supplementary Figure S6. Sequence logo comparison of matched and unmatched reference TSS promoters.** Sequence logo analysis comparing promoters derived from matched reference TSSs and promoters derived from the remaining unmatched reference TSSs under the 100-bp tolerance setting. Promoter sequences were separated into the −10 element, −35 element, and spacer regions. Compared with unmatched TSS promoters, matched TSS promoters showed stronger and more conserved promoter sequence patterns, particularly in the −10 element and the G-rich region near the downstream end of the spacer.

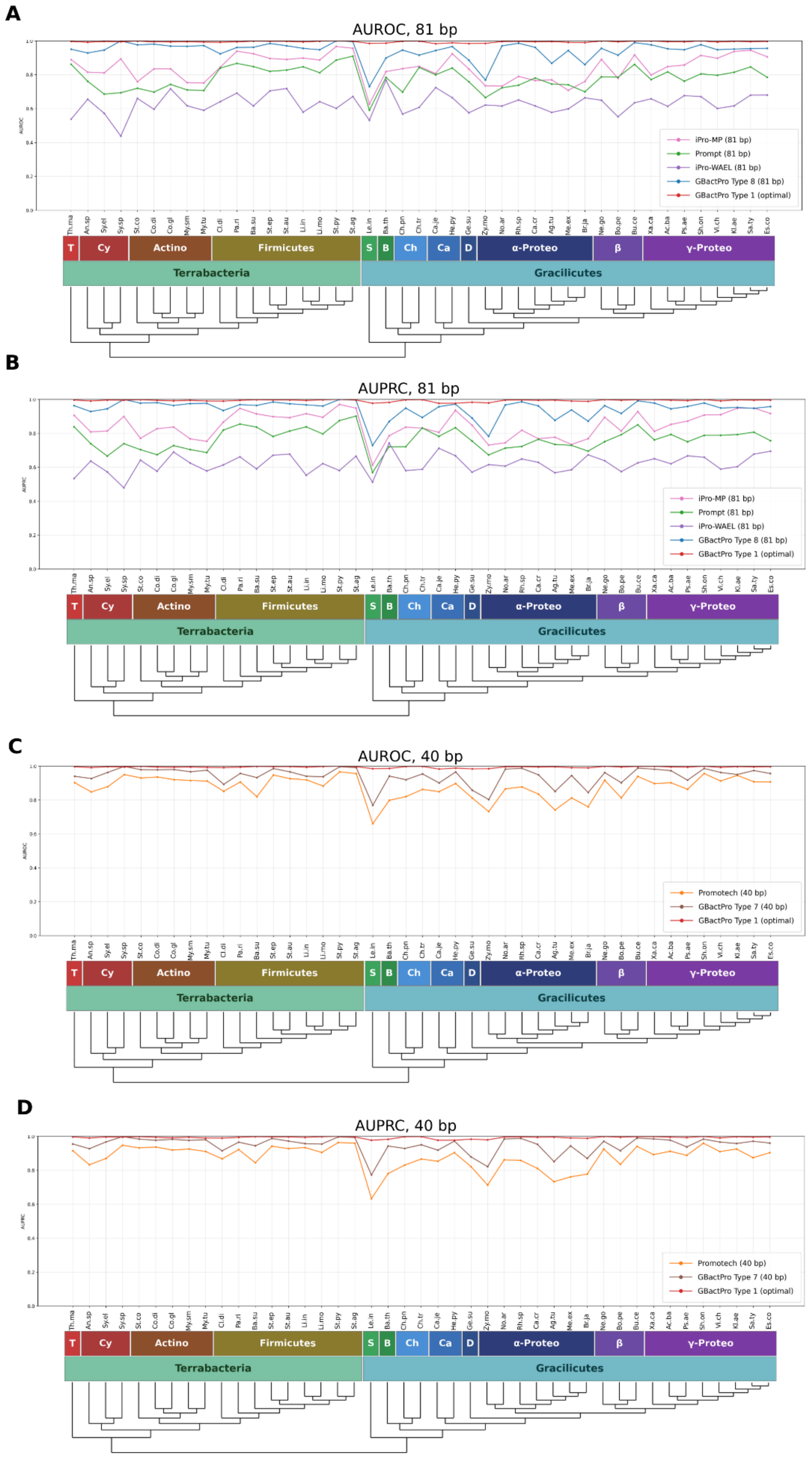
A
AUROC, 81 bp
AUROC
iPro-MP (81 bp)
Prompt (81 bp)
iPro-WAEL (81 bp)
GBactPro Type 8 (81 bp)
GBactPro Type 1 (optimal)
Th.ma An.sp Sy.el Sy.sp St.co Co.di Co.gl My.sm My.tu Cl.di Pa.ri Ba.su St.ep St.au Li.in Li.mo St.py St.ag Le.in Ba.th Ch.pn Ch.tr Ca.je He.py Ge.su Zy.mo No.ar Rh.sp Ca.cr Ag.tu Me.ex Br.ja Ne.go Bo.pe Bu.ce Xa.ca Ac.ba Ps.ae Sh.on Vi.ch Kl.ae Sa.ty Es.co
T Cy Actino Firmicutes S B Ch Ca D α-Proteo β γ-Proteo
Terrabacteria
Gracilicutes
B
AUPRC, 81 bp
AUPRC
iPro-MP (81 bp)
Prompt (81 bp)
iPro-WAEL (81 bp)
GBactPro Type 8 (81 bp)
GBactPro Type 1 (optimal)
C
AUROC, 40 bp
AUROC
Promotech (40 bp)
GBactPro Type 7 (40 bp)
GBactPro Type 1 (optimal)
D
AUPRC, 40 bp
AUPRC
Promotech (40 bp)
GBactPro Type 7 (40 bp)
GBactPro Type 1 (optimal)

**Supplementary Figure S7. Comparison of GBactPro with benchmark promoter prediction models across 43 bacterial species.** (A) Per-species AUROC under the matched 81-bp input setting. The jointly trained 43-species GBactPro model using the Type 8 design was compared with iPro-MP, Prompt, and iPro-WAEL using the same held-out test sequences; the optimal Type 1 GBactPro model is shown for reference. (B) Corresponding AUPRC comparison under the 81-bp setting. (C) Per-species AUROC under the matched 40-bp input setting. The jointly trained 43-species GBactPro model using the Type 7 design was compared with Promotech using the same held-out test sequences; the optimal Type 1 model is shown for reference. (D) Corresponding AUPRC comparison under the 40-bp setting.

### Supplementary Tables

**Supplementary Table S1. Training and testing dataset composition across 43 bacterial species.** For each species, the complete dataset was divided into 75% training and 25% testing sets. Training data were constructed with a 1:10 positive-to-negative ratio to reflect the rarity of promoters in genomic sequence, whereas testing data used a balanced 1:1 ratio to support consistent sequence-level evaluation across species.

| | Training | | Testing | |
|---|---|---|---|---|
| **Species** | **pos** | **neg** | **pos** | **neg** |
| Thermotoga maritima | 412 | 4120 | 138 | 138 |
| Anabaena sp. | 2966 | 29660 | 989 | 989 |
| Synechococcus elongatus | 1821 | 18210 | 608 | 608 |
| Synechocystis sp. | 127 | 1270 | 43 | 43 |
| Streptomyces coelicolor | 762 | 7620 | 254 | 254 |
| Corynebacterium diphtheriae | 901 | 9010 | 301 | 301 |
| Corynebacterium glutamicum | 1610 | 16100 | 537 | 537 |
| Mycolicibacterium smegmatis | 2282 | 22820 | 761 | 761 |
| Mycobacterium tuberculosis | 1333 | 13330 | 445 | 445 |
| Clostridioides difficile | 966 | 9660 | 322 | 322 |
| Paenibacillus riograndensis | 951 | 9510 | 318 | 318 |
| Bacillus subtilis | 450 | 4500 | 150 | 150 |
| Staphylococcus epidermidis | 904 | 9040 | 302 | 302 |
| Staphylococcus aureus | 1350 | 13500 | 450 | 450 |
| Listeria innocua | 683 | 6830 | 228 | 228 |
| Listeria monocytogenes | 809 | 8090 | 270 | 270 |
| Streptococcus pyogenes | 512 | 5120 | 171 | 171 |
| Streptococcus agalactiae | 663 | 6630 | 222 | 222 |
| Leptospira interrogans | 2148 | 21480 | 716 | 716 |
| Bacteroides thetaiotaomicron | 1212 | 12120 | 404 | 404 |
| Chlamydia trachomatis | 261 | 2610 | 88 | 88 |
| Chlamydia pneumoniae | 286 | 2860 | 96 | 96 |
| Campylobacter jejuni | 506 | 5060 | 169 | 169 |
| Helicobacter pylori | 537 | 5370 | 179 | 179 |
| Geobacter sulfurreducens | 656 | 6560 | 219 | 219 |
| Zymomonas mobilis | 2403 | 24030 | 802 | 802 |
| Novosphingobium aromaticivorans | 708 | 7080 | 236 | 236 |
| Rhodobacter sphaeroides | 815 | 8150 | 272 | 272 |
| Caulobacter crescentus | 1082 | 10820 | 361 | 361 |
| Agrobacterium tumefaciens | 351 | 3510 | 117 | 117 |
| Methylorubrum extorquens | 1522 | 15220 | 508 | 508 |
| Bradyrhizobium japonicum | 3338 | 33380 | 1113 | 1113 |

| Species | Training | | Testing | |
|---|---|---|---|---|
| | **pos** | **neg** | **pos** | **neg** |
| Neisseria gonorrhoeae | 689 | 6890 | 230 | 230 |
| Bordetella pertussis | 444 | 4440 | 149 | 149 |
| Burkholderia cenocepacia | 814 | 8140 | 272 | 272 |
| Xanthomonas campestris | 1158 | 11580 | 387 | 387 |
| Acinetobacter baumannii | 648 | 6480 | 216 | 216 |
| Pseudomonas aeruginosa | 1587 | 15870 | 530 | 530 |
| Shewanella oneidensis | 1351 | 13510 | 451 | 451 |
| Vibrio cholerae | 962 | 9620 | 321 | 321 |
| Klebsiella aerogenes | 351 | 3510 | 117 | 117 |
| Salmonella enterica Typhimurium | 840 | 8400 | 280 | 280 |
| Escherichia coli | 1398 | 13980 | 467 | 467 |

**Supplementary Table S2. Hyperparameters of the CNN–BiLSTM architecture.** Layer-by-layer specification of the CNN–BiLSTM model, including filter counts, kernel sizes, BiLSTM units, dropout rate, and training hyperparameters.

| Category | Hyperparameter | Value |
|---|---|---|
| Input | Encoding | One-hot (A,T,C,G → 4 channels) |
| Architecture | Conv block 1 | 2× Conv1D (32 filters, k=3 & k=5), BN, MaxPool, Dropout 0.5 |
| | Conv block 2 | 2× Conv1D (64 filters, k=3 & k=5), BN, MaxPool, Dropout 0.5 |
| | BiLSTM layer | 1024 units, ReLU activation |
| | Hidden dense | 1024 units, ReLU activation |
| | Output | 1 unit, sigmoid activation |
| Training | Optimizer | Adam (α = 0.001) |
| | Loss function | Binary cross-entropy |
| | Batch size | 32 |

**Supplementary Table S3. Summary of model architecture, input design, training scope, and benchmark evaluation for GBactPro and comparator models.** Architecture, input-length specification, training-data scope, publication source, and evaluation design are summarized for GBactPro and the four benchmark tools used in the matched-length comparison. All tools were evaluated on the same held-out test sequences.

| Model | Type | Architecture / feature design | Input length | Training scope | Publication |
|---|---|---|---|---|---|
| GBactPro | Deep-learning hybrid | 1D-CNN blocks + BiLSTM | 27–31 bp | 43 species | — |
| iPro-MP | Transformer | Pre-trained DNABERT-6 + MLP classifier | 81 bp | Fine-tuned on E. coli | Genome Biology, 2025 |
| Prompt | Neural-network ensemble | MLP ensemble with Tanh + dropout | 81 bp | Trained on E. coli | Interdiscip. Sci., 2024 |
| iPro-WAEL | Ensemble | Weighted ensemble of RF + Word2Vec-CNN | 81 bp | Trained on E. coli | Nucleic Acids Research, 2022 |
| Promotech | Machine learning | Random forest on one-hot sequence features | 40 bp | 9 species | Genome Biology, 2021 |

**Supplementary Table S4.** Mean cross-species AUROC and rankings across 43 bacterial species. For each species, the mean training AUROC was calculated across the corresponding row of the 43 × 43 cross-species evaluation matrix, whereas the mean testing AUROC was calculated across the corresponding column. Species were ranked separately based on their mean training and testing AUROC values.

| | Training | | Testing | |
|---|---|---|---|---|
| **Species** | **Mean AUROC** | **AUROC Rank** | **Mean AUROC** | **AUROC Rank** |
| Thermotoga maritima | 0.956 | 40 | 0.978 | 23 |
| Anabaena sp. | 0.993 | 1 | 0.961 | 38 |
| Synechococcus elongatus | 0.990 | 3 | 0.963 | 35 |
| Synechocystis sp. | 0.882 | 43 | 0.967 | 33 |
| Streptomyces coelicolor | 0.979 | 22 | 0.974 | 26 |
| Corynebacterium diphtheriae | 0.984 | 12 | 0.971 | 30 |
| Corynebacterium glutamicum | 0.989 | 4 | 0.971 | 29 |
| Mycolicibacterium smegmatis | 0.984 | 11 | 0.971 | 31 |
| Mycobacterium tuberculosis | 0.983 | 14 | 0.964 | 34 |
| Clostridioides difficile | 0.978 | 24 | 0.974 | 27 |
| Paenibacillus riograndensis | 0.980 | 20 | 0.977 | 25 |
| Bacillus subtilis | 0.972 | 33 | 0.986 | 8 |
| Staphylococcus epidermidis | 0.974 | 31 | 0.985 | 9 |
| Staphylococcus aureus | 0.987 | 7 | 0.986 | 7 |
| Listeria innocua | 0.975 | 29 | 0.981 | 18 |
| Listeria monocytogenes | 0.981 | 19 | 0.982 | 15 |
| Streptococcus pyogenes | 0.961 | 36 | 0.994 | 1 |
| Streptococcus agalactiae | 0.972 | 34 | 0.990 | 2 |
| Leptospira interrogans | 0.988 | 6 | 0.929 | 43 |
| Bacteroides thetaiotaomicron | 0.979 | 21 | 0.962 | 36 |
| Chlamydia trachomatis | 0.961 | 37 | 0.983 | 12 |
| Chlamydia pneumoniae | 0.957 | 39 | 0.982 | 17 |
| Campylobacter jejuni | 0.974 | 30 | 0.959 | 39 |
| Helicobacter pylori | 0.977 | 25 | 0.977 | 24 |
| Geobacter sulfurreducens | 0.964 | 35 | 0.954 | 40 |
| Zymomonas mobilis | 0.986 | 8 | 0.945 | 42 |
| Novosphingobium aromaticivorans | 0.953 | 41 | 0.979 | 21 |
| Rhodobacter sphaeroides | 0.959 | 38 | 0.988 | 6 |
| Caulobacter crescentus | 0.976 | 27 | 0.982 | 16 |
| Agrobacterium tumefaciens | 0.953 | 42 | 0.984 | 10 |
| Methylorubrum extorquens | 0.982 | 18 | 0.962 | 37 |
| Bradyrhizobium japonicum | 0.990 | 2 | 0.954 | 41 |
| Neisseria gonorrhoeae | 0.978 | 23 | 0.988 | 4 |
| Bordetella pertussis | 0.973 | 32 | 0.979 | 22 |
| Burkholderia cenocepacia | 0.975 | 28 | 0.988 | 5 |

| | Training | | Testing | |
|---|---|---|---|---|
| **Species** | **Mean AUROC** | **AUROC Rank** | **Mean AUROC** | **AUROC Rank** |
| Xanthomonas campestris | 0.984 | 10 | 0.980 | 20 |
| Acinetobacter baumannii | 0.983 | 17 | 0.973 | 28 |
| Pseudomonas aeruginosa | 0.983 | 15 | 0.969 | 32 |
| Shewanella oneidensis | 0.983 | 16 | 0.988 | 3 |
| Vibrio cholerae | 0.983 | 13 | 0.984 | 11 |
| Klebsiella aerogenes | 0.976 | 26 | 0.982 | 14 |
| Salmonella enterica Typhimurium | 0.985 | 9 | 0.981 | 19 |
| Escherichia coli | 0.989 | 5 | 0.983 | 13 |
